# PHARAOH: A collaborative crowdsourcing platform for PHenotyping And Regional Analysis Of Histology

**DOI:** 10.1101/2024.03.20.585977

**Authors:** Kevin Faust, Min Li Chen, Parsa Babaei Zadeh, Dimitrios Oreopoulos, Alberto J. Leon, Evelyn Rose Kamski-Hennekam, Marly Mikhail, Xianpi Duan, Xianzhao Duan, Mugeng Liu, Narges Ahangari, Raul Cotau, Vincent Francis Castillo, Nikfar Nikzad, Richard J. Sugden, Patrick Murphy, Susan Done, Safiyh S. Aljohani, Philippe Echelard, Kiran Jakate, Yazeed Alwelaie, Mohammed J. Alyousef, Noor Said Alsafwani, Assem Saleh Alrumeh, Rola Saleeb, Maxime Richer, Lidiane Vieira Marins, George M. Yousef, Phedias Diamandis

## Abstract

Deep learning has proven to be capable of automating key aspects of histopathologic analysis, but its continual reliance on large expert-annotated training datasets hinders widespread adoption. Here, we present an online collaborative portal that streamlines tissue image annotation to promote the development and sharing of custom computer vision models for PHenotyping And Regional Analysis Of Histology (PHARAOH; https://www.pathologyreports.ai/). PHARAOH uses a weakly supervised active learning framework whereby patch-level image features are leveraged to organize large swaths of tissue into morphologically-uniform clusters for batched human annotation. By providing cluster-level labels on only a handful of cases, we show how custom PHARAOH models can be developed and used to guide the quantification of cellular features that correlate with molecular, pathologic and patient outcome data. Both custom model design and feature extraction pipelines are amenable to crowdsourcing making PHARAOH a fully scalable systems-level solution for the systematic expansion and cataloging of computational pathology applications.

---

Deep learning has the potential to help automate and objectify many manual and subjective aspects of histomorphologic analysis^1^. With the growing availability of digital Hematoxylin and Eosin (H&E)-stained tissue whole slide images (WSIs), machines can now leverage massive volumes of labeled image data to guide feature engineering in an entirely automated and data-driven manner. Despite this exciting prospect, real-world utility and broad adoption of deep learning in pathology has been challenged by the high input requirements of expert-annotated data for each context-specific application. To solve this gap, recent approaches have explored using weakly-supervised multi-instance learning to assign patch-level tissue labels using existing WSI-level clinical annotations^2^. While these approaches have shown that laborious manual annotations can be potentially bypassed, the models’ continued reliance on massive data volumes to achieve good performance (e.g. ∼10,000 WSIs/application) limits generalizability^2^. There is therefore a need for more practical hybrid approaches where both humans and machines contribute to learning, as these may provide a more favourable balance between automation and data efficiency. Such “active” learning paradigms^3,4^ may help to more efficiently produce the full diversity and scale of training data necessary to generate a comprehensive toolbox of context-specific convolutional neural networks (CNNs) for diverse histomorphologic analysis applications.

We recently developed a computational pipeline that takes advantage of deep learning feature vectors (DLFVs), generated in a CNN’s final global pooling layer, to serve as “histomorphologic fingerprints” of individual pathology images^5^. By clustering associated image patches using these signatures, we found that we could organize WSI, from a wide diversity of tissue types, into regional partitions that show relative cytoarchitectural uniformity^6^ (**Fig 1Ai**). Importantly, these proposed tissue regions correlate with expert annotations, immunohistochemical readouts and even subtle intra-tumoral differences in molecular profiles^7^. For image annotation, this DLFV-based clustering approach therefore has the potential to serve as the foundation of an active weakly-supervised learning approach in which a system only needs to query experts for sparse cluster-level labeling of grouped image patches for custom model training^3,4^. By combinatorically coupling these different tissue segmentation models with a complementary set of tissue-agnostic cell phenotyping readouts, we build a framework for an expansive community-driven encyclopedia of computational pathology tools for PHenotyping and Regional Analysis Of Histology (PHARAOH; (https://www.pathologyreports.ai/) (**Fig 1Aii-iv**).

**Figure 1.**
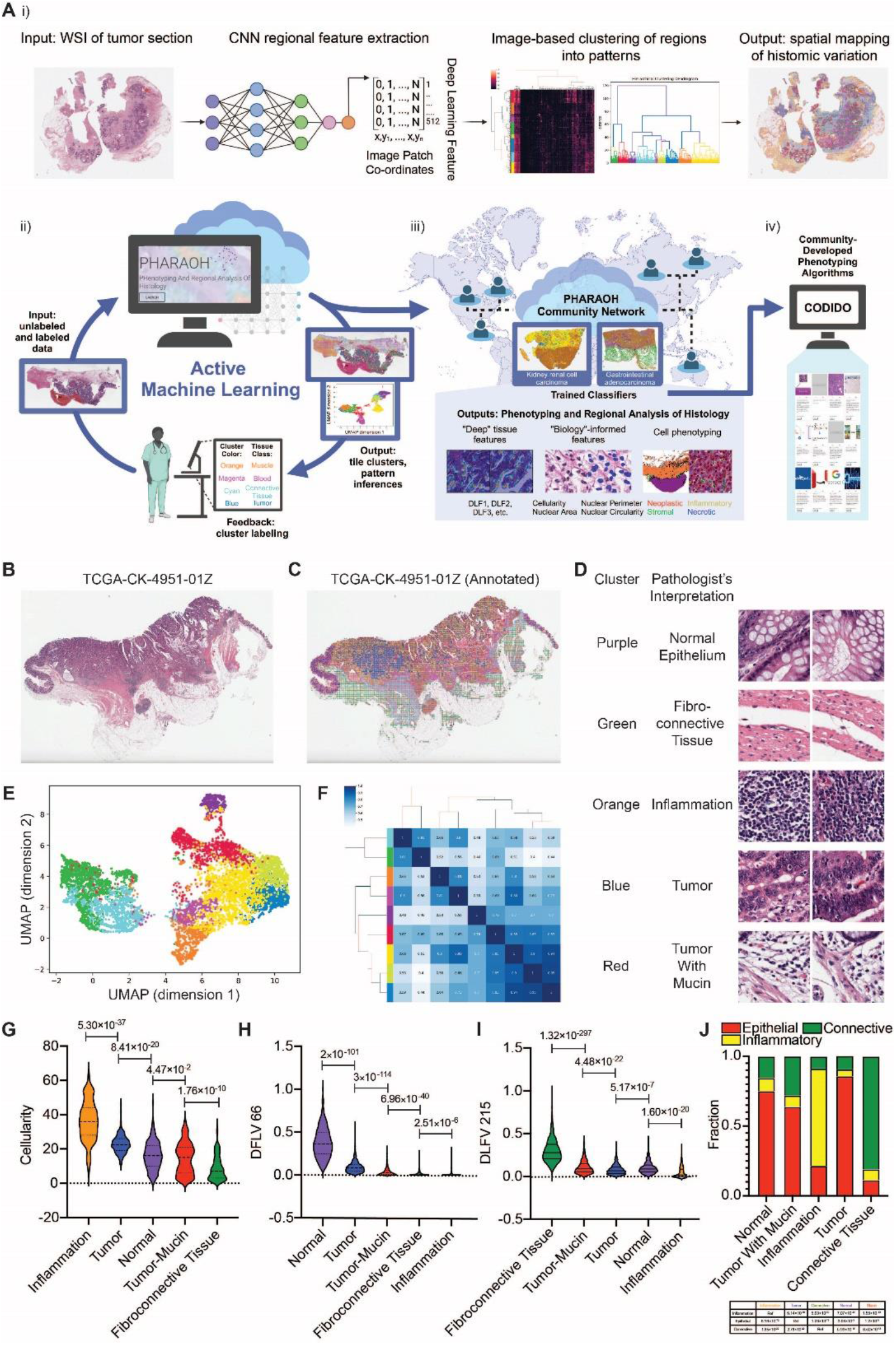
Deep learning feature-based clustering partitions histomorphologically-complex tissue WSIs into distinct and relatively uniform image patch subgroups. (**A i)**) Workflow highlighting image feature-based clustering and mapping of tissue patterns across entire WSIs. Briefly, a pre-trained CNN is used as a feature extractor and the generated tile-level DLFVs are clustered and mapped back onto the WSI. (**A ii – iv)**) Cartoon schematic of the PHARAOH workflow (https://www.pathologyreports.ai/). ii). unlabeled WSIs are uploaded by histologists to the online portal. Users receive image feature-based clustering maps and other visualization tools to help decipher unsupervised histomorphologic tile groupings. Users provide cluster-level annotations which are aggregated across multiple WSIs and used to finetune custom CNN models. This iterative active learning process is repeated to further refine accuracy and desired outputs. iii). Once developed, trained classifiers are made available to users across the entire PHARAOH network. In addition to tissue segmentation, various regional histomic (DLFs) and cell-based phenotyping outputs are provided to serve as biomarkers of disease and relevant clinical outcomes (e.g. Tumor infiltrating lymphocytes). iv) In addition to core PHARAOH outputs, users can export segmented regions of interest and carry out custom image analyses using other third-party tools on companion platforms (CODIDO; codido.co). (**B-C**) Demonstrative input (WSI) (panel B) and output (tissue heterogeneity map) (panel C) images of a sample colorectal adenocarcinoma from The Cancer Genome Atlas (TCGA). Scale bar: 6 mm. (**D**) Representative image patches highlighting stereotypical morphology from different partitions. (**E-F**) The relative degree of histomorphology similarities and differences align with cluster positioning on dimensionality reduction plots (e.g. UMAP) (panel E) and Pairwise Pearson correlation coefficients (r) (panel F) of the DFLVs of proposed partitions. (**G-I**) Violin plots highlighting quantitative cellularity (Mask R-CNN-based quantification) (panel G), epithelial (DLF66) (panel H), and fibrosis (DLF215) (panel I) marker differences between defined regions. The violin plot shows minimum, first quartile, median, third quartile, and maximum. Counts represent nuclear instances per 67,488 μm^2^. (**J**) Regional cell composition differences (generated using HoVer-Net outputs).

Here, we use pathologic, molecular and clinical outcome data to assess if human annotations applied to PHARAOH-proposed clusters, each comprising ∼10^2^-10^3^ related image patches, can be used to fine-tune CNNs for custom applications in computational pathology. To organize histological information spanning entire tissue sections into meaningful partitions, PHARAOH divides submitted WSIs into non-overlapping image patches (0.017-0.07 mm^2^; user-defined size) and uses their individual DLFV signatures to organize tiles a empirically optimized solution of nine image clusters^7^ (**Fig 1B-C**). Importantly, when tile partitions are projected back onto the WSI, clusters often show non-random spatial distributions that closely mirror morphologic regions of uniformity that can be easily recognized and intuitively labeled by pathologists (**Fig 1D**). The four major tissue types/patterns annotated in this demonstrative case (fibroconnective tissue in green/cyan clusters; normal epithelium in purple cluster; inflammation in orange cluster and tumor in blue/lime clusters) are also supported by the positional relationships of patches on uniform manifold approximation and projection (UMAP) and pairwise Pearson correlation coefficients (r) of regional DLFVs (**Fig 1E-F**).

The downstream outputs of PHARAOH also help objectively benchmark the uniformity of the delineated histologically tissue regions. For example, PHARAOH routinely performs nuclear segmentation/classification and basic morphometric analyses (e.g. cellularity and nuclear surface area) in a subset of tiles derived from each tissue region. Indeed, as highlight in this example, cellularity counts are, as expected, significantly higher in compact lymphocyte-rich tissue regions as compared to areas containing larger neoplastic and non-neoplastic epithelial cells, as well as the more paucicellar connective tissue compartments (**Fig 1G**). Previously-reported individual deep learning features (DLFs)^5^ are also standard PHARAOH outputs and closely correlate with epithelial- (DLF66), fiber- (DLF215) and mucin- (DLF382) rich regions, allowing users to have additional supporting and objective metrics of interpretability of different cluster compositions (**Fig1H-I & Supplementary Fig S1**). Finally, regional nuclear segmentation and classification using the HoVerNet-PanNuke^8,9^ model can also provide support for an increased relative number of tumor, inflammatory and/or stromal cells within respective WSI partitions (**Fig 1J**). Overall, this framework of regional phenotyping with interpretable features provides a dynamic approach for arranging large swaths of tissue into relatively uniform subgroups that show concordance with annotations provided by human experts as well as a number of corroborating quantitative and objective morphometric outputs.

To demonstrate the suitability of our workflow for the nimble development of clinically-meaningful histopathologic applications, we examined the relationship between the levels of intratumor lymphocytes (TILs), inferred from H&E images, and median overall survival time in patients with cutaneous melanoma. First, we performed automated region delineation in a handful of WSI samples (n=7) from the cutaneous melanoma cohort of the Cancer Genome Atlas (TCGA-SKCM) (**Supplementary Fig S2**). On average, we found labeling of tissue regions delineated by the PHARAOH workflow took ∼1-2 minutes/WSI, and generated a total of 23,211 annotated images spanning 8 unified classes (e.g. tumor and surrounding tissue types) (**Supplementary Table 1**). This weakly-labeled dataset was then used to fine-tune a VGG19 CNN, which could then be used to segment tumor regions in the remaining TCGA-SKCM cases via tile-based classification (PHARAOH model ID: e5dad8db). Specifically, for this experiment, we used the cell phenotyping readouts provided by HoVer-Net/PanNuke to quantify immune cells in up to 200 representative tumor-enriched tiles per case (probability score: >90% tumor) (**Fig 2A**). All together, after filtering out low-quality WSI, we applied this tandem model to compute a sample-level TIL score for the remaining 385 patients from TCGA-SKCM cohort, defined as the log-transformed mean value of lymphocyte counts across the set of sampled tiles classified as tumor (**Supplementary Table 2**). Molecular metadata in TCGA supported the validity of our approach as the computed TIL score correlated with the RNA-based Lymphocyte Infiltrating Signature Score^10^, (R^2^ = 0.18, *p*<2.2e^-16^) (**Fig 2C**). Moreover, partitioning the TCGA-SKCM cohort into cases with either high or low TIL scores (with respect to the median score) showed a significant survival difference (high=109 months, low = 66 months, log rank test, *p* = 0.0016) (**Fig 2D**), which aligns with the previously-reported results from RNA-based TIL inferences in RNA^11^ and other more manually curated computer vision TIL models^12^. Together, this demonstrates how the annotation of image-feature-based clusters, generated from a handful of cases submitted to PHARAOH, can be utilized to develop custom biologically-informative histologic biomarkers of clinical outcomes by imposing only modest computational and data curation requirements on users.

**Figure 2.**
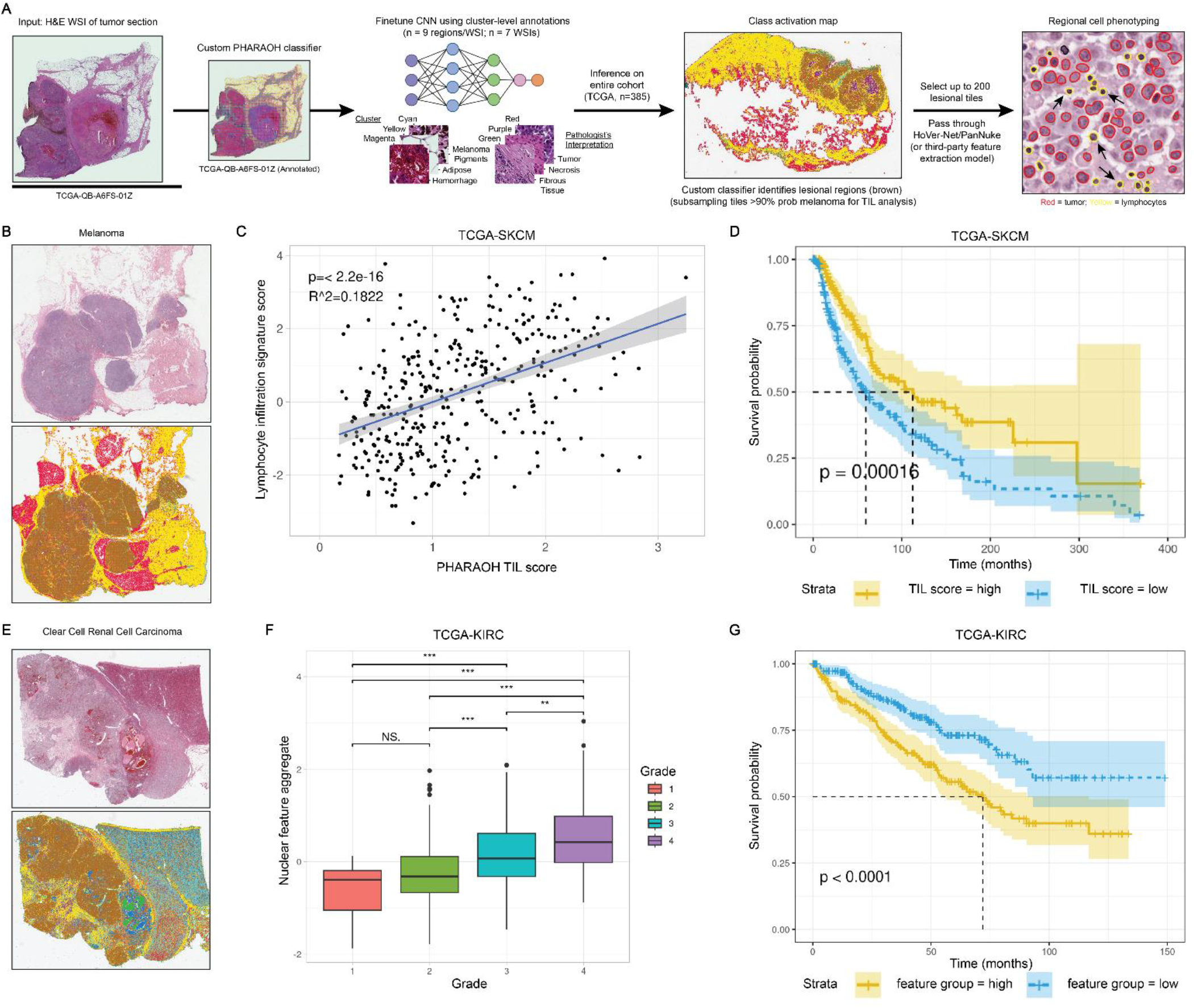
Tumor segmentation and regional nuclear phenotyping, guided by sparse cluster-level annotations, predicts outcomes in cutaneous melanoma and clear cell renal cell carcinoma. (**A**) Weakly-supervised annotation pipeline for custom CNN training and TIL inferencing. (**B**) Sample case input and class activation map (CAM) output of a representative case from the TCGA-SKCM cohort. Custom region of interest (melanoma) is shown in brown, while adipose and fibroconnective tissue are shown in red and yellow, respectively. (**C**) Scatter plot of case-level correlation between the PHARAOH-based TIL quantification and the RNA-based Lymphocyte infiltration signature score across the TCGA-SKCM cohort. P-value generated by linear fit model. (**D**) Kaplan-Meier survival curves for the TCGA-SKCM cohort split into “high” and “low” PHARAOH-TIL scores based on the overall cohort’s median value. (**E**) Sample case input and CAM output of a representative case from the TCGA-KIRC cohort. Custom region of interest (RCC) is shown in brown while normal renal parenchyma and fibroconnective tissue are shown in cyan and yellow, respectively. (**F**) Box plot showing aggregate nuclear feature values for TCGA-KIRC cases, separated by their reported TCGA nuclear grades. (**G**) Kaplan-Meier survival curves for the TCGA-KIRC cohort split into “high” and “low” aggregate nuclear feature score based on the overall cohort’s median value.

To further promote the design of custom applications, users can leverage the PHARAOH models to delineate tumor-containing areas of their submitted WSIs, and then export the representative set of custom image tiles of interest for independent downstream analysis. To facilitate this in a collaborative manner, we developed a companion platform to help crowdsourced and host executable machine learning tools (CODIDO; https://www.codido.co/). As a proof-of-concept, we investigated the degree of correlation between the grading annotations of the TCGA clear cell renal cell carcinoma cohort (TCGA-KIRC) and relevant nuclear morphometric features computed externally with CellProfiler^13^. The aforementioned weakly supervised learning framework was used to develop a tumor segmentation model for clear cell renal cell carcinoma (ccRCC) with cluster-level annotations for 27 cases from the TCGA-KIRC cohort. A total of 81,768 annotated images spanning 9 tissue classes were generated (**Supplementary Table 3 & Supplementary Fig S3**), and later used to fine-tune a custom RCC model (PHARAOH model ID: 82829b22). We then used this PHARAOH model to extract a representative set of up to 200 tumor-enriched tiles (probability score: >90% tumor) from each of the remaining 475 TCGA-KIRC cases. These tiles were exported and analyzed using the CellProfiler-based “Nuclear Feature Extractor” workflow (https://www.codido.co/marketplace/browse?RepoInfo%5Bquery%5D=feature%20extractor). Briefly, each case’s set of tiles underwent nuclear segmentation using HoVer-Net and then each nuclear object was subjected to morphometric analysis with CellProfiler (v4.0.5) resulting in 78 nuclear features. To approximate the Fuhrman ccRCC grading system, reported in TCGA-KIRC samples, we created a composite score using the average of the normalized values of the following three relevant features: (i) nuclear atypia (inverse AreaShape_FormFactor), (ii) nuclear size (AreaShape_Perimeter) and (iii) nuclear staining heterogeneity as a marker of prominent nucleoli (Intensity_MADIntensity_Grayscale) (**Supplementary Table 4**). Importantly, the benchmarking of this aggregate score with the reported TCGA grading assignments of the remaining cases showed a significant correlation (**Fig 2F**). Similarly, partitioning the TCGA-KIRC cohort into cases with either high or low aggregate nuclear scores (with respect to the median cohort value) showed a significant survival difference (log rank test, *p* = 0.0001) (**Fig 2G**). In addition to further supporting the ability of PHARAOH to serve as a platform for the development of reliable tumor segmentation models, these results also demonstrate how the platform can work cooperatively with external tools to combinatorically expand the repertoire of clinically-meaningful outputs that can be collaboratively generated by users.

Deep learning has proven to be capable of addressing many of the challenges surrounding human subjectivity and automation in histomorphologic analysis. However, the inherent context-specific nature of developing robust models, including the development of domain-specific training image datasets, has made implementation and translational efforts difficult. To address these barriers in a scalable manner, we developed PHARAOH, a generalizable weakly-supervised workflow to streamline the development of custom computational pathology models. Importantly, models developed on PHARAOH are shared and accessible to the research community via the open web portal (pathologyreports.ai). Notably, the cluster-level labels PHARAOH uses significantly reduces the time investment from expert annotators, who have demanding clinical schedules, and can be carried out remotely without the need for any specialized hardware/software. Indeed, we collectively used this pipeline to virtually catalog a diverse set of models that span several international institutions and among collaborators with no previous relationships.

By coupling the contextual flexibility of CNNs for accurate tissue segmentation (region of interest selection) with the generalizability of different cell-based phenotyping tools (for feature extraction), we show how clinically-meaningful and interpretable tissue biomarkers can be quickly developed through the annotation of only a handful of relevant cases. Importantly, the development and hosting of both PHARAOH classifiers and external third-party feature extractors are highly amenable to community crowdsourcing and sharing by expert histologists and computer scientists, respectively. An added benefit of making these digital pathology models publicly available is that it facilitates peer-based benchmarking of developed histologic biomarkers across various institutions and diverse patient populations. Together, we believe PHARAOH provides a versatile infrastructure to accelerate the scaling and adoption of computer vision in histopathologic analysis.

## Acknowledgments

This work is supported by the Canadian Institute of Health Research and the Princess Margaret Cancer Foundation.

## Author contributions

K.F. and P.D. conceived the idea and approach. K.F. & A.L. developed the computational pipelines. K.F., M.C., A.L. and D.O. developed the weakly-supervised custom training approach. K.F, M.C., A.L., E.K.H, and P.B.Z. analyzed and interpreted data outputs. D.O., M.M, K.J., P.M., P.B.Z., L.M., M.R., N.S.A., R.C., R.S., V.F.C., N.N., Y.A., M.A.Y., G.Y., A.S.A., S.D., S.S.S.A., and P.E. provided annotations and designed and tested custom classifiers using the developed approach. K.F. developed the online portal. K.F., R.S., X.D., X.D, M.L, developed the CODIDO workflow and relevant models. K.F., A.L. and P.D. wrote the manuscript, with input from all other authors. P.D. supervised the work.

## Methods

### Deep image feature-based decomposition of WSIs into patch clusters with relatively uniform histomorphology

Tissue partitions are generated in PHARAOH using an unsupervised image feature-based clustering workflow implemented in python (https://pypi.org/project/havoc-clustering/), as previously described^6,7^. Briefly, WSIs are first tiled into individual 0.066-0.27 mm^2^ image patches (patch width: 129 μm (256 pixels) to 258 μm (512 pixels)) respectively. We found that this magnification effectively separates a variety of tissue types and tumor sub-patterns and provides a favourable balance between capturing both individual cellular differences (e.g. nuclear features) and more advanced secondary structures. Although various tile sizes were used during the initial tests, for simplicity and consistency, we have set the current default on PHARAOH to 0.066 mm^2^ (512 x 512 pixel, 20x apparent magnification). We found the results generated from this patch size, maintained easily recognizable spatial patterns in tissue makeup while reducing tile-to-tile variability and not compromising the computational time of the workflow.

Histomorphologic signatures for individual tiles are represented by averaging the “deep learning feature” (DLF) values extracted from the final global average pooling layer of a previously fine-tuned version of the VGG19 CNN that we trained, with transfer learning, on a diverse set of nearly 1 million pathologist-annotated image patches spanning over 70 distinct tissue classes that were extracted from over 1,000 brain tumors samples^5^. We refer to these 512 feature representations as the “deep learning feature vector” (DLFV). We previously showed that individual DLFs are activated by specific histomorphologic patterns (e.g. fibrosis, epithelium, and mucin), allowing them to drive the clustering of image patches with relatively similar morphologies^5^. Tile-level DLFVs are scaled feature-wise and are then hierarchically clustered into 9 clusters using Ward’s Method^5^. Empirically, this solution tends to ensure slight over-clustering of distinct (sub)regions while maximizing the production of relatively uniform histomorphological subgroups of images that are readily identifiable by expert reviewers. To help further qualitatively and quantitatively visualize inter-cluster relationships, PHARAOH also produces tile-level UMAP projections and pair-wise Pearson correlation coefficients of each region’s average DLFVs.

### Weakly-supervised annotation and tissue-specific CNN fine-tuning

The PHARAOH workflow generates tissue-specific CNN classifiers by using expert-annotated histologically homogeneous clusters of images. Briefly, WSIs undergo deep image-based decomposition, as described above, to form 7-12 partitions, usually comprising ∼10^3^-10^4^ image patches per cluster (256px, 20x apparent magnification). To facilitate the annotation process, PHARAOH also provides a companion set of interpretable features at the region level. Using Mask R-CNN^14^ for nuclear segmentation and Detectron2 for morphometric analysis, the workflow calculates regional average values for cell counts, nuclear surface area and circularity, as described previously^15^. Additionally, “preliminary labels” and probability scores from exiting models are provided for each region to pre-populate commonly found histological entities and allow users to focus their annotations on the classes that lacked high-confidence labeling. For each WSI, users are provided with 2 files for annotation: a (i) colour-coded cluster WSI map (Format: .jpg, Size: ∼3-5 mb) and a (ii) spread sheet (Format: .cvs, Size: ∼2 Kb) containing the list of tissue regions along with basic regional metrics (cellularity, nuclear surface area, # tiles/region) and “preliminary labels” to allow users to provide custom batch-level annotations. The annotation files from multiple WSIs are merged, and by evaluating a summary file, users can confirm whether all the tissue classes have reached a sufficient number of annotated tiles for CNN fine-tuning.

The annotated collection of image patches is then used to fine-tune a CNN. The image set is filtered to remove images with over 40% blank space, and the remaining images are then partitioned into training and validation sets (ratio: 85:15). The final, fully-connected layers of the VGG19 CNN are removed and replaced with a global average pooling, single fully-connected layer, and then, these final two convolutional layer blocks of the network are retrained using the user-annotated images as previously described^16^. The process is carried out using the Keras framework with a Tensorflow backend and powered by an NVIDIA RTX 3090 graphics processing unit (GPU). Users are free to retrain and refine models as they see fit.

### Nuclear segmentation and classification with HoVer-Net/PanNuke

To provide estimates of the cellular composition of the tissue images, PARAOH performs nuclear segmentation and classification using the HoVer-Net/PanNuke model (TIAToolbox implementation^17^), which produces counts for the following cell types: neoplastic epithelial, inflammatory, connective, necrotic, and non-neoplastic epithelial. Although this model was trained with 512 x 512 pixel tiles and 40x apparent magnification, we can also apply this model to a WSI scanned at an apparent magnification of 20x by producing tiles of 256 x 256 pixel and scaling them to pseudo-40x magnification, 512 x 512 pixel, using bicubic pixel interpolation (‘vips resize’ command). To manage the level of utilization of computational resources, nuclear segmentation and classification is carried out on a sample of up to 200 tiles from each region of interest.

As part of the effort to generate interpretable readouts for the regions produced by automated tissue segmentation, PHARAOH runs HoVer-Net/PanNuke in tiles sampled from all the resulting regions. In Fig 1, we merged the non-neoplastic and neoplastic epithelial categories and omitted the necrotic class as we found that these classes had low specificity in this context.

After a tissue classification model has been established, HoVer-Net/PanNuke can be run in the tiles that have been classified as lesional with high degree of confidence (>0.9 probability) to provide a readout of the immune status of the tumor areas. A total of 476 WSIs (.svs files) from the TCGA-SKCM study^18^ were downloaded from the GDC Data Portal using the GDC Data Transfer Tool Client v1.6.1. The associated clinical information was obtained from cBioportal’s dataset download section (https://www.cbioportal.org/datasets) by selecting the “TCGA, Firehose legacy” data release of the TCGA-SKCM study. We developed a training cohort of labeled image patches (256 x 256 pixel, 20x apparent magnification) using the PHARAOH workflow as described above. Specifically, we selected 7 representative WSIs containing melanoma and other commonly-encountered tissue types including normal skin, lymph nodes and intestinal mucosa. Each WSI was passed through our image feature-based clustering workflow to delineate 12 clusters of tiles that captured 8 tissue classes (**Supplementary Fig S2** and **Supplementary Table 1**). The annotated tiles were used to fine-tune a pre-trained VGG19 neural network. The remaining WSIs of the TCGA-SKCM were analyzed with the newly developed melanoma classifier, and a representative sample from each WSI of up to 200 tiles (bicubic pixel interpolation for 512 x 512 pixel, pseudo-40x magnification) was analyzed with the HoVer-Net/PanNuke model. For this analysis of the TCGA-SKCM cohort, we made the assumption that the resulting inflammatory cell counts provided by the model are equivalent to TIL counts. A region-level TIL score was defined using the following formula:

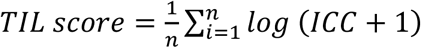

In this formula, TIL score is the mean value of the log-transformed “inflammatory cell count” (ICC) of each selected tile (n = up to 200 available tiles/region) computed by HoVer-Net/PanNuke.

### Nuclear segmentation and morphometric analysis in RCC

To investigate the level of alignment between nuclear morphometric features and the pathology grading (Fuhrman grade) in RCC, the PHARAOH weakly-supervised workflow was used to develop a RCC classifier with 9 tissue types using images from TCGA-KIRC (n=27) and additional non-lesional tiles from TCGA-KIRP (n=2) (**Supplementary Table 3**), and using 512 x 512 pixel tiles with 20x apparent magnification. The classifier was then used to analyze the remaining WSIs from the TCGA-KIRC cohort to delineate their tumor-containing regions, followed by sampling of up to 200 lesional tiles from each WSI. HoVer-Net/Kumar^8^, an image segmentation architecture trained to detect cell nuclei in histology images, was applied to each set of lesional tiles. Then, a CellProfiler pipeline was used to perform morphometric analyses in the extracted cell nuclei, including the image analysis modules MeasureObjectSizeShape, MeasureImageAreaOccupied, MeasureObjectIntensity, MeasureObjectIntensityDistribution (with maximum radius set to 100) and MeasureGranularity. A total of 78 morphometric features were generated, and their values averaged for each WSI. We selected three morphometric features that recapitulate known pathologic hallmarks of RCC, namely AreaShape_FormFactor (nuclear atypia), AreaShape_Perimeter (nuclear size) and Intensity_MADIntensity_Grayscale (nuclear stain heterogeneity). To facilitate the statistical analyses, an aggregate score was calculated as the average of the normalized values for AreaShape_Perimeter and Intensity_MADIntensity_Grayscale, and using the inverse values for AreaShape_FormFactor. A Docker image with CellProfiler was used as base component of the workflow, to which HoVer-Net/Kumar and application-specific scripts were added (https://github.com/duanxianpi/Nuclei-Feature-Extraction). This workflow can be run either in a GPU-enabled system locally or online in PHARAOH’s companion platform CODIDO (https://www.codido.co/; “Nuclear Feature Extractor”).

