## Supplementary Figure S1 for "PHARAOH: A collaborative crowdsourcing platform for PHenotyping And Regional Analysis Of Histology"

**Supplementary Figure S1.** Feature Activation Mapping (FAM) highlights tissue patterns enriched within specific compartments of WSIs. A-B Representative feature activation maps (FAMs) of DLF66 (panel A) and DLF215 (panel B) highlighting non-overlapping epithelial and connective tissue-rich regions, respectively. C. H&E and respective FAMs of the highest and lowest activating tiles within the mucinous tumor partition (RED) for DLF66, DLF215 and DLF382. Even within a specific partition, the tiles with the highest values of the aforementioned features show an enrichment of recognizable epithelial, fibrous and mucinous tissue. FAMs show appropriate coordinates of activation of the specific features. Together with the average partition-level values in Figure 1, these provide support for the uniform partitioning of human-recognizable tissue patterns by the image-feature clustering workflow to facilitate intuitive batch annotations and interpretability.

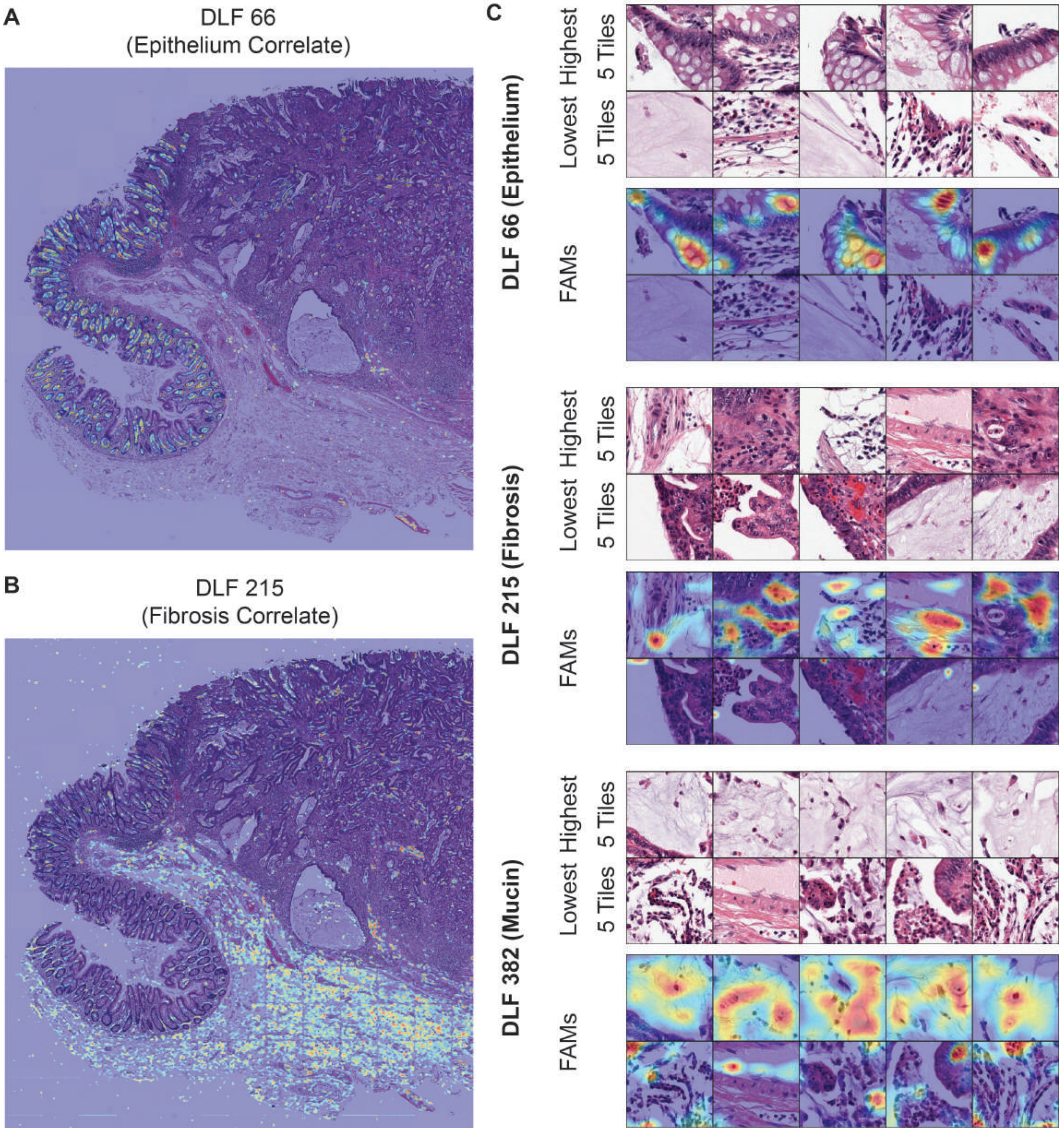
