## Supplementary Figure S2 for "PHARAOH: A collaborative crowdsourcing platform for PHenotyping And Regional Analysis Of Histology"

**Supplementary Figure S2.** Representative examples of automated region segmentation and annotated tiles from different classes that were used to train the PHARAOH melanoma classifier.

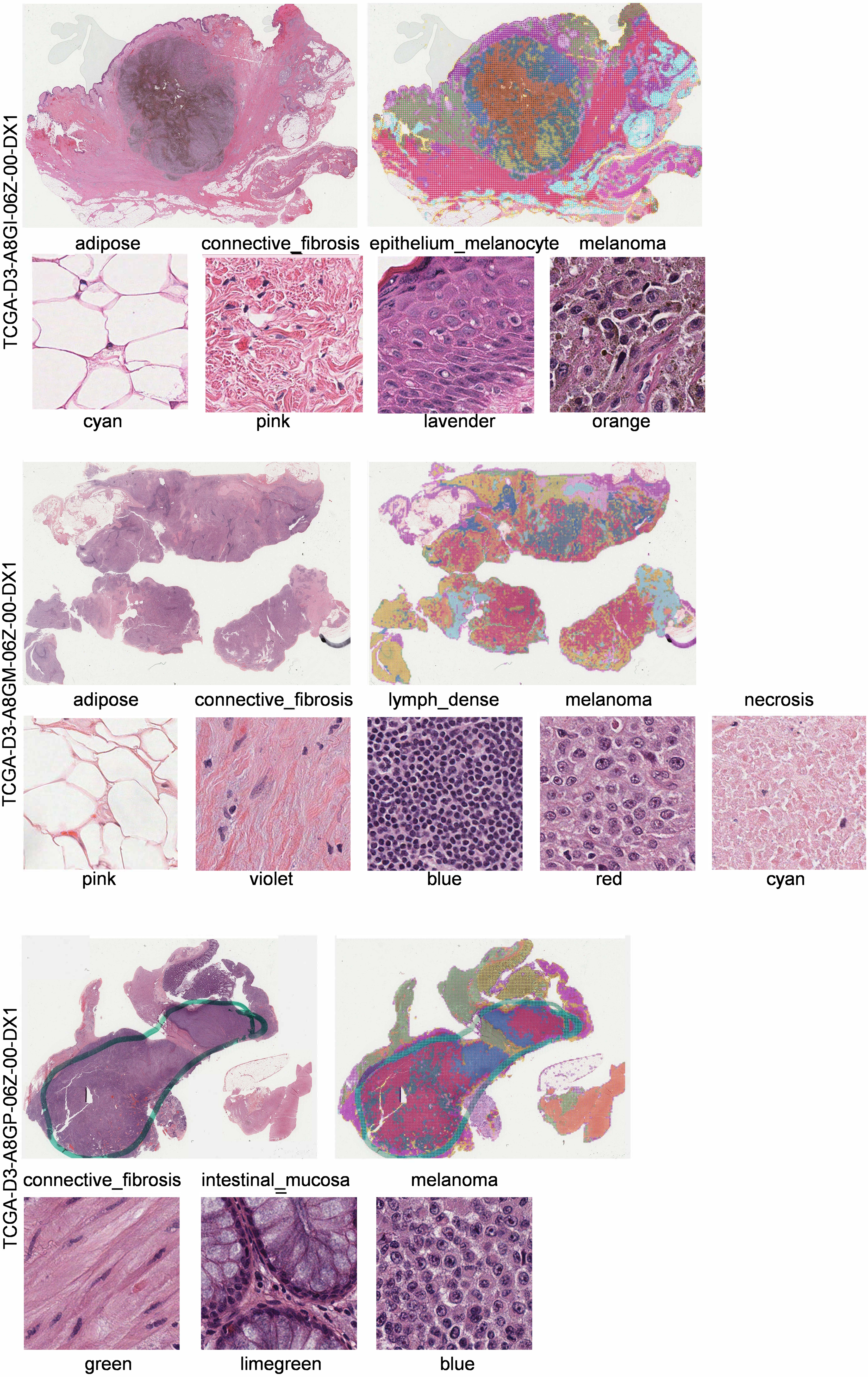
