## Supplementary Figure S3 for "PHARAOH: A collaborative crowdsourcing platform for PHenotyping And Regional Analysis Of Histology"

**Supplementary Figure S3.** Representative examples of automated region segmentation and annotated tiles from different classes that were used to train the PHARAOH clear cell renal cell carcinoma (ccRCC) classifier.

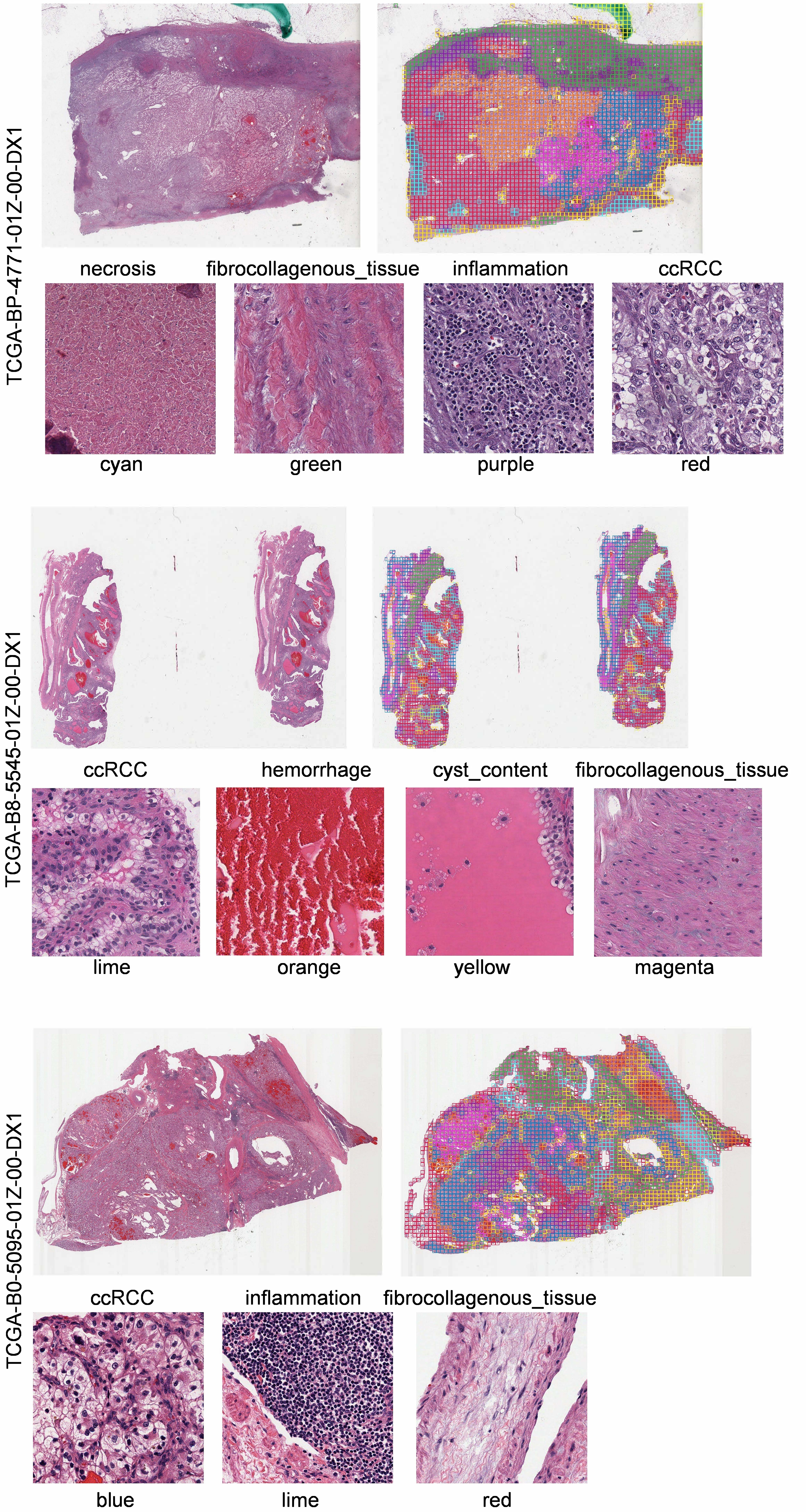
